# Genome Mining and Characterization of a Heme-Dependent Enzyme Catalyzing Intermolecular Nitrogen–Nitrogen Bond Formation in Hydrazinosuccinic Acid Biosynthesis

**DOI:** 10.1101/2025.06.26.661531

**Authors:** Jingkun Shi, Zhijie Zhao, Jian Yang, Ziyang Cheng, Hu Li, Yu Liu, Guiyun Zhao, Miaolian Wu, Yi-Ling Du

## Abstract

Nitrogen–nitrogen (N–N) bond-forming enzymes are rare but play vital roles in both primary and secondary metabolism. Guided by a nitric oxide synthase (NOS)-based genome mining strategy, we report the discovery and characterization of a new heme-dependent enzyme system that catalyzes intermolecular N–N bond formation. Using both in vivo and in vitro reconstitution approaches, we demonstrated that a protein complex, comprising a heme enzyme and a 2[4Fe–4S] ferredoxin partner, mediates the coupling of the α-amine group of L-aspartate with inorganic nitrogen oxide species, such as nitrite or nitric oxide, to generate hydrazinosuccinic acid, a key biosynthetic precursor in several natural product pathways. Structural modeling and site-directed mutagenesis suggest a plausible catalytic mechanism involving the formation of a reactive nitrogen intermediate, potentially a heme-bound nitrene species. These findings reveal a new family of N–N bond-forming biocatalysts that leverage inorganic nitrogen sources, offering valuable tools for genome mining and the synthetic biology.

## Introduction

Nitrogen is the fourth most abundant element in biological systems. Although nitrogen gas (N2) makes up nearly 80% of the Earth’s atmosphere, only a limited number of organisms can convert atmospheric nitrogen into biologically accessible forms [1,2]. Nitrogen availability in ecosystems is regulated by a series of microbial transformation processes, including nitrogen fixation, nitrification, denitrification, and ammonification. Traditionally, it has been believed that nitrogen assimilation occurs mainly through the incorporation of ammonia (NH_4_+) into organic nitrogen-containing compounds. However, recent studies have begun to reveal that other inorganic nitrogen species in the nitrogen cycle—such as nitric oxide (NO), and nitrite (NO_2_−), hydrazine (N_2_H_4_)—can also serve as direct biosynthetic precursors, and be incorporated by microorganisms into the molecular frameworks of various bioactive secondary metabolites, including thaxtomin A, rufomycin, 8-azaguanine and alanosine (Figure 1a) [3–6]. In parallel, new biochemical pathways responsible for generating these inorganic nitrogen species have also begun to emerge. For instance, in the biosynthetic pathways of cremeomycin and several other bacterial natural products, an enzyme pair (CreD/CreE) was shown to catalyze the sequential oxidation of the α-amine of aspartic acid, ultimately releasing it as nitrite [7]. Additionally, a distinct pathway leading to N2H4 production has been implicated in the biosynthesis of the antibiotic albofungin [8].

**Figure 1.**
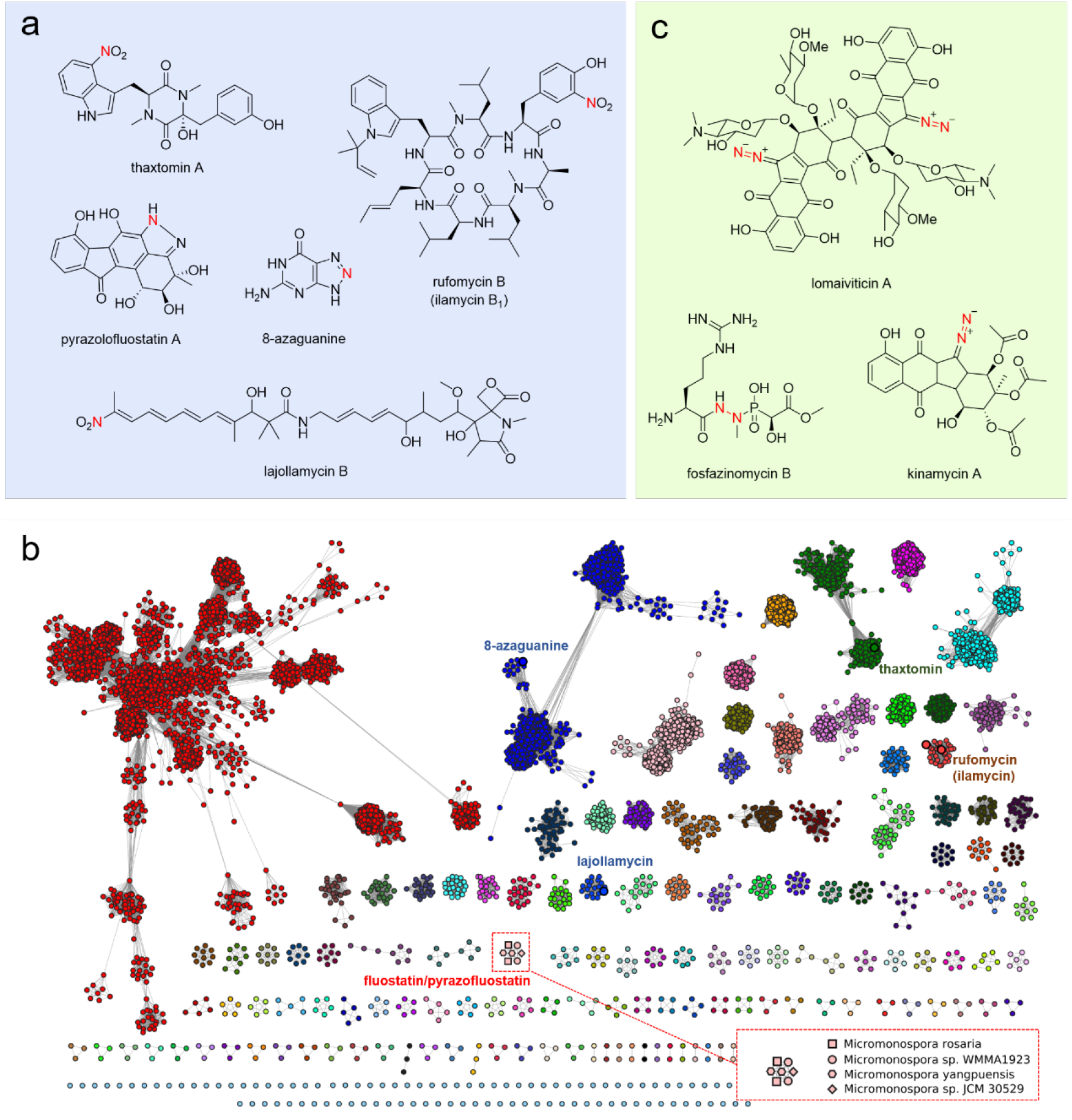
Examples of microbial natural products using inorganic nitrogen species as a direct nitrogen donor and. (a) Compounds with nitric oxide as a building block. (b) Sequence similarity network (SSN) analysis of bacterial nitric oxide synthases (NOSs). Note: NOSs present in the BGCs of known natural products are displayed next to the corresponding nodes. The cluster of NOSs for further investigation is boxed within red dash line, and the strain information are also displayed. (c) Compounds with nitrite as a building block.

The ability of microorganisms to both produce and utilize inorganic nitrogen species as direct nitrogen donors for specialized metabolism suggests the existence of previously unrecognized biochemical routes for nitrogen assimilation. Indeed, in the biosynthetic pathways of compounds such as thaxtomin A and rufomycins, cytochrome P450 enzymes have been shown to catalyze aromatic nitration using nitric oxide as the nitrogen source [3]. More recently, a class of ATP-dependent enzymes that utilize nitrite has been identified to catalyze N–N bond formation via energy-driven coupling reactions, yielding diazo or triazene-containing products [9–11].

Given that genes involved in related biosynthetic functions are often co-localized in gene clusters, we hypothesized that genes responsible for production of inorganic nitrogen species—such as NO, NO_2_−, and N_2_H_4_—could serve as genetic markers for the discovery of novel enzymes that directly employ these species in nitrogen-incorporation chemistry. Supporting this hypothesis, we recently used the bacterial nitric oxide synthase (NOS) gene as a genomic probe and identified a novel cytochrome P450-like enzyme capable of olefin nitration in lajollamyicin biosynthesis [12]. Here, using a combination of NOS-guided genome mining, in vivo reconstitution, and in vitro biochemical characterization, we characterized a heme-dependent enzyme that catalyzes intermolecular N-N bond formation between aspartic acid (L-Asp) and either nitrite or nitric oxide. This reaction yields hydrazinosuccinic acid (**1**), a metabolic precursor in the biosynthetic pathway of various bioactive metabolites. These findings not only expand the current enzymatic toolkit for N–N bond formation but also provide a valuable biocatalyst for the synthesis of nitrogen-containing compounds in synthetic biology.

## Results

### Discovery of potential NO-utilizing enzymes by genome mining using bacterial nitric oxide synthase as a probe

For genome mining, we use the protein sequence of the nitric oxide synthase (NOS) PtnF from the 8-azaguanine biosynthetic gene cluster as a query [5]. A BLASTP search was performed against the NCBI non-redundant (nr) protein database (maximum target sequences = 50,000; all other parameters set to default). From the results, the top 10,000 protein sequences were selected and submitted to the EFI-EST (Enzyme Function Initiative-Enzyme Similarity Tool) web server for sequence similarity network (SSN) analysis using an E-value cutoff of 150 [13]. Guided by this NOS-based genome mining strategy, we observed that a group of putative NOS genes from several *Micromonospora* strains formed a distinct cluster in our sequence similarity network (SSN) analysis (Figure 1c). Previous studies using heterologous expression have demonstrated that, the NOS-containing genomic regions, originating from an environmental DNA sample and the genome of *Micromonospora rosaria*, encode the biosynthetic gene cluster (BGC) for fluostatins and pyrazolofluostatins [14–16]. This finding suggested that one of the nitrogen atoms in the structure of pyrazolofluostatin may be derived from NO or a related nitrogen species, and that the BGC likely also encodes additional enzyme(s) involved in the utilization of NO or NO-derived species (Figure 1a). Notably, the fluostatins/pyrazolofluostatins BGC shares a conserved sub-cluster of five genes (*flsT-S-N3-N4-U2*) with the BGCs of several other bacterial natural products containing a nitrogen–nitrogen (N–N) bond, including kinamycin, fosfazinomycin and lomaiviticins [17–20] (Figure 1c nd Figure 2a). Previous investigation of this conversed gene cassette from the BGCs of fosfazinomycin (*fzmR-Q-P-O-N*) and kinamycin (*kinJ-K-L-M-N*) have revealed its involvement in the biosynthesis of glutamylhydrazine, a common intermediate in these pathways (Figure S1) [21]. Among these genes, four of them (FzmQRNO or KinNMLK) have been shown to catalyze sequential conversion of hydrazinosuccinic acid (**1**) to glutamylhydrazine.

**Figure 2.**
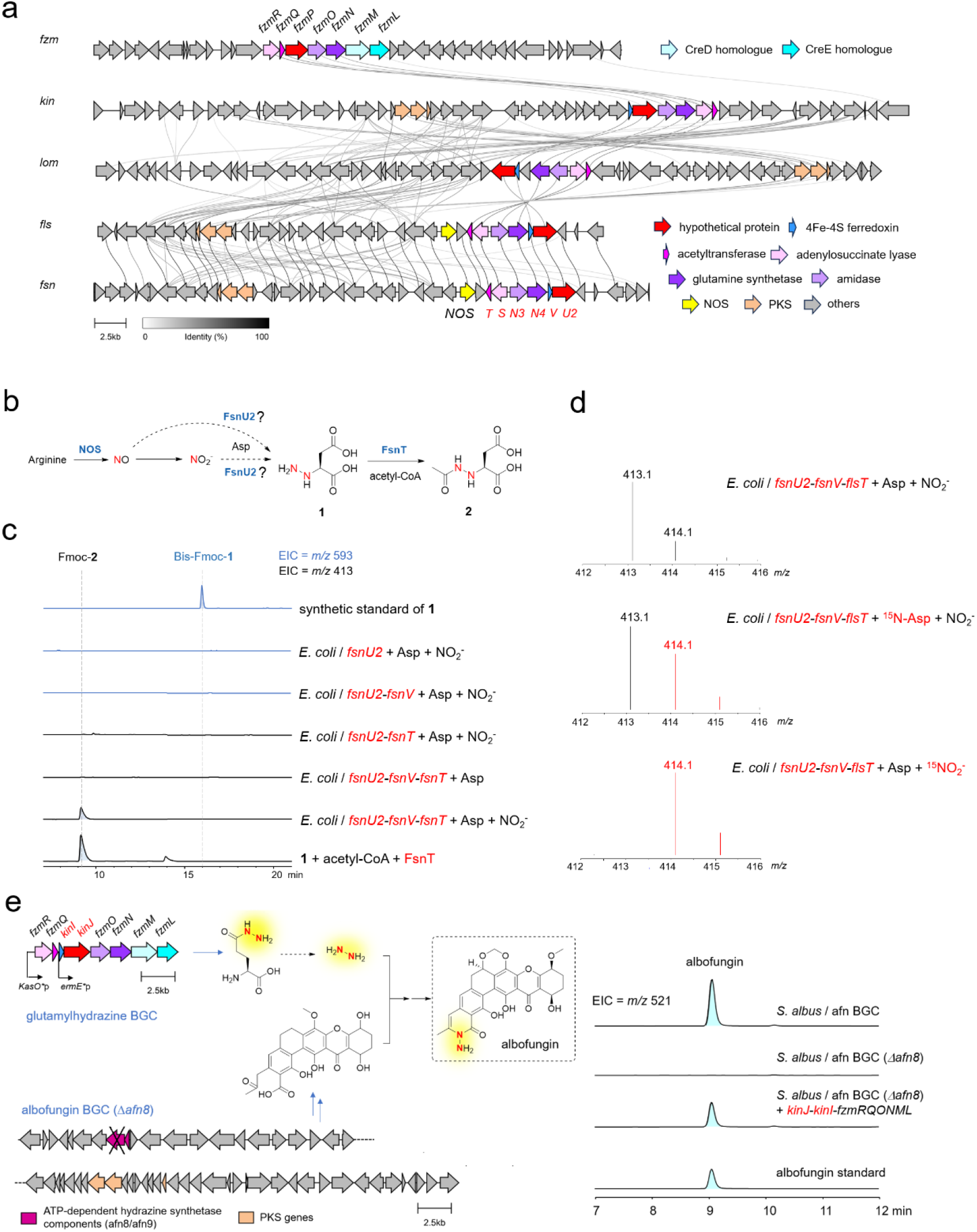
In vivo reconstitution identifies a protein pair for hydrazinosuccinic acid biosynthesis. (a) Comparison of biosynthetic gene clusters harboring conserved gene cassette involved in hydrazinosuccinic acid biosynthesis. (b) Proposed reaction route for hydrazinosuccinic acid formation. (c) In vivo reconstitution in *Escherichia coli* identifies the essential role of *fsnU2* and *fsnV* in hydrazinosuccinic acid biosynthesis. (d) LC-MS analysis of the isotopic pattern of compound Fmoc-**2** after the producer strain was fed with ^15^NO_2_^−^ or ^15^N-Asp. (e) In vivo reconstitution in *Streptomyces* using albofungin biosynthetic system as a reporter for hydrazine production. Note; *afn8* is a key component of ATP-dependent hydrazine synthetase in albofungin BGC, and mutation in this gene accumulates a polyketide intermediate that can undergoes enzymatic hydrazine incorporation.

While hydrazinosuccinic acid itself has been proposed to originate from an N–N coupling reaction between L-aspartate (L-Asp) and nitrite, based on isotopic labeling experiments and the presence of nitrite-generating genes (*creDE* homologs) within the corresponding BGCs, the enzyme(s) responsible for catalyzing this unusual N–N bond formation remain unidentified [19,22]. Given that the function of the remaining gene (*fzmP*/*kinJ*) in the conserved gene cassette for glutamylhydrazine biosynthesis is still unknown, it was thus proposed that FzmP/KinJ may catalyze the N–N bond formation by linking nitrite (NO_2_−) to the amino group of L-Asp to form **1**. This hypothesis, however, has not yet been experimentally confirmed. In contrast to the BGCs of kanamycin and fosfazinomycin, which contain *creDE* genes for NO_2_− production, the fluostatin/pyrazolofluostatin BGC lacks *creDE* homologs but instead encodes a NOS gene (Figure 2a). This raises intriguing questions about the N–N bond forming reaction to afford compound **1**, which is likely mediated by the hypothetical protein including FzmP, KinJ, and their counterparts such as FlsU2 and FsnU2, from *Micromonospora rosaria* and *Micromonospora* sp. JCM 35029 (Figure 1c, 2a and 2b). We next set out to characterize this potential new N-N bond forming enzyme family.

### In vivo reconstitution identifies a protein pair for hydrazinosuccinic acid biosynthesis

Because the biosynthesis of hydrazinosuccinic acid (**1**) likely involves an unusual condensation between Asp and nitrite or nitric oxide, a reaction that is rarely observed in nature, we first employed an in vivo reconstitution approach to gain insights into this reaction. The gene *fsnU2* was cloned from the strain *Micromonospora* sp. JCM 35029, and then expressed in *Escherichia coli* (Figure 1c). The culture of the resulting strain

*E. coli*/*fsnU2* was then supplemented with L-Asp and nitrite. However, LC-MS analysis of the culture supernatant did not reveal any mass signal corresponding to **1**, using synthetic **1** as a reference standard (Figure 2c and Figure S2). We noticed that *fsnU2* (and its homologs from the kinamycin and lomaiviticin BGCs) is positioned adjacent to a gene encoding a putative [4Fe–4S] ferredoxin (*fsnV*) (Figure 2a). Given that the formation of **1** via N-N coupling between NO/NO_2_^−^ and L-Asp potentially involves redox chemistry, we thus co-expressed *fsnU2* with *flsV* in *E. coli*. Additionally, to stabilize the potential product **1** under in vivo conditions, we also included the *N*-acetyltransferase *fsnT*, which can convert **1** into a more stable product, *N*-acetly-**1** (**2**) (Figure 2b). The supernatants of strains *E. coli*/*fsnU2*-*fsnV* and *E. coli*/*fsnU2*-*fsnV-fsnT* were then similarly analyzed following pre-column derivatization with Fmoc-chloride. While compound **1** remained undetected in strain *E. coli*/*fsnU2*-*fsnV*, a mass signal corresponding to Fmoc-**2** (*m*/*z* 413, [M+H]^+^) was successfully observed in strain *E. coli*/*fsnU2*-*fsnV-fsnT* (Figure 2c). The production of this compound was dependent on both FsnT and the presence of nitrite, and its retention time matched that of Fmoc-**2**, which was synthesized enzymatically by incubating FsnT with acetyl-CoA and **1**, followed by Fmoc-Cl derivatization (Figure S3). Furthermore, LC-MS analysis of this compound from strain *E. coli*/*fsnU2*-*fsnV-fsnT* cultures fed with either ^15^N-labeled L-Asp or ^15^NO_2_- revealed the enrichment of +1 isotopic peaks (Figure 2d). Taken together, these results support that FsnU2 and FsnV function cooperatively to catalyze the intermolecular N-N bond formation between L-Asp and a nitrigen oxide (NO_x_) species, leading to the production of **1**. To further test whether the homologs of FsnU2 and FsnV from other BGCs are also capable of catalyzing the same reaction, we generated strain *E. coli*/*kinJ-kinI-fsnT* and found that this strain indeed produced **2** (Figure S4).

To further explore whether the FsnU2/FsnV or KinJ/KinI enzyme pairs are also sufficient to mediate hydrazine formation in vivo within an actinomycete host, we employed the albofungin biosynthetic system as a functional reporter (Figure 2e). This system is based on our previous finding that free hydrazine can be enzymatically incorporated into the characteristic N-amino amide moiety of albofungin [8]. We speculated that glutamylhydrazine, once produced, might undergo hydrolysis in vivo to release of free hydrazine, which could then be utilized by the albofungin biosynthetic pathway. To test this, we constructed a refactored BGC containing *kinJ*/*kinI* and other glutamylhydrazine biosynthetic genes, and introduced it into a *Streptomyces albus* host horboring an albofungin BGC that lacks its native hydrazine producing genes. LC-MS analysis revealed the the expression of this *kinJ*/*kinI*-containing cassette successfully restored albofungin production in the engineered strain (Figure 2e). These results support the conclusion that KinJ/KinI, and by extention FsnU2/FsnV and their homologs, could represent a new biosynthetic strategy for N-N bond formation in nature.

### In vitro characterization of the protein pair responsible for N-N bond formation

Next, we sought to reconstitute the FsnU2/FsnV-catalyzed reaction in vitro. His6-tagged recombinant FsnU2 and FsnV proteins were overexpressed and purified using nickel affinity chromatography. The purified FsnU2 appeared brownish in color and exhibited characteristic UV-visible absorption spectra (with γmax at 418 nm) consistent with the presence of a heme cofactor. Upon addition of dithionite, the Soret band shifted to 429 nm (Figure 3a). Further analysis confirmed that FsnU2 contains a noncovalently bound b-type heme as its cofactor (Figure S5). Collectively, these results supporting the classification of FsnU2 as a heme-binding enzyme.

**Figure 3.**
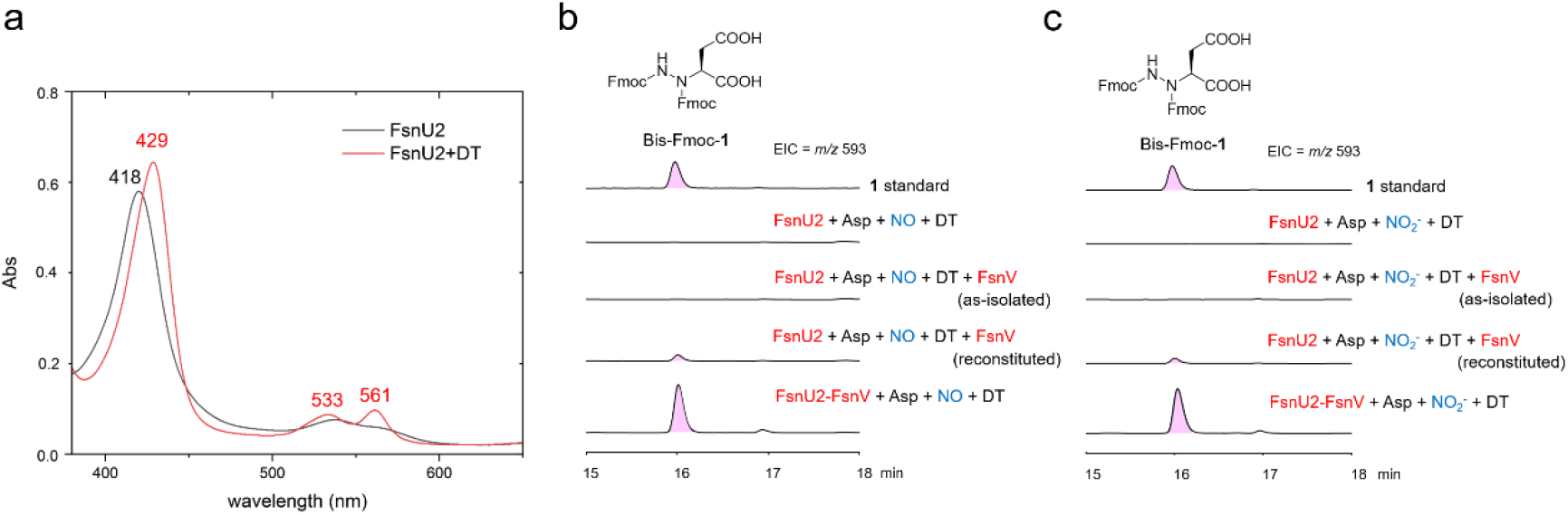
In vitro characterization of the protein pair FsnU2/FsnV that catalyzes N-N bond formation. (a) UV-Vis spectra of as-isolated FsnU2 (oxidized) and dithionite (DT)-reduced FsnU2. (b) and (c) In vitro assays of using nitric oxide (b) or nitrite (c) as nitrogen donor. The extracted ion chromatograph (EIC = *m*/*z* of 593, [M+H]^+^ for bis-Fmoc-**1**) from LC–MS analysis of the in vitro reaction mixtures were displayed. All the reactions were conducted under anaerobic conditions. Note: for the reactions with NO, the NO donor Diethylamine NONOate was used.

Given previous finding on heme-dependent piperazate synthase—an enzyme that catalyzes intramolecular N-N bonds via direct substrate coordination to the heme iron, we hypothesized that the FsnU2-catalyzed reaction proceeds through a similar mechanism [23]. However, in the case of FsnU2, the reaction likely involves one or more reductive steps acting on nitrite or nitric oxide prior to N–N bond formation. To test this, we carried out in vitro biochemical assays by incubating L-Asp and either nitrite or nitric oxide (NO) with FsnU2 and as-isolated FsnV in the presence of dithionite (Figure 3b). Since the [4Fe-4S] cluster of FsnV is prone to degradation during purification, we also reconstituted its iron-sulfur cluster inside a glovebox and used the reconstituted protein for enzyme assays (Figures 3a-3b and Figure S6). All the in vitro biochemical assays for FsnU2 and FsnV were performed under anaerobic conditions.

LC-MS analysis of the reaction mixtures showed that a small amount of product **1** was generated in assays containing reconstituted FsnV, regardless of whether nitrite or NO was used as the substrate (Figure 3b and 3c). This result highlights the necessity of active FsnV for the FsnU2-mediated transformation. Several commonly used redox partners for P450 enzymes were also tested for their ability to support the FsnU2-catalyzed reaction, including seFdx/seFdR from *Synechococcus elongatus*, CamB (putidaredoxin)/CamA (putidaredoxin reductase) from *Pseudomonas putida*, an Fdx/anFdR from *Anabaena* PCC7119, and RhFRED from *Rhodococcus* [24–26]. However, none of these redox systems supported the production of compound **1** in vitro. Interestingly, we observed that the His6-tagged FsnU2 protein purified from the *E. coli* strain co-expressing fsnU2 and untagged fsnV resulted in co-purification of a protein band corresponding to FsnV, suggesting the formation of a FsnU2–FsnV protein complex (Figure S7). Notably, when this co-purified FsnU2–FsnV complex was directly used in the in vitro assay, a substantially higher amount of **1** was produced compared to assays using separately purified FsnU2 and FsnV (Figure 3b and 3c). These findings suggest that complex formation between FsnU2 and FsnV may enhance the stability and/or catalytic efficiency of one or both proteins. We also evaluated hydroxylamine (NH_2_OH) as an alternative substrate for the FsnU2–FsnV complex, However, no production of **1** was detected under the conditions tested (Figure S8). Although the specific in vivo redox partner(s) for the FsnU2-FsnV complex in the *E. coli* or *Streptomyces* remain unidentified, the combined in vivo and in vitro data described above, supports the conclusion that FsnU2 functions in concert with FsnV to catalyze N–N bond formation.

### Mechanistic insights into hydrazinosuccinic acid biosynthesis

Our efforts to crystallize either FsnU2 alone or the FsnU2–FsnV complex have thus far been unsuccessful. Consequently, we employed AlphaFold to predict the structures of FsnU2, FsnV, and the FsnU2–FsnV complex (Figure 4 and S9-S11) [27]. In the predicted structure of FsnU2 bound to heme, the heme cofactor is coordinated by His532, a residue conserved among FsnU2 homologs, including KinJ and FzmP (Figure 4a-4b and Figure S12). In the structural model of FsnV, two sets of conserved cysteine residues (Cys39, Cys42, Cys45, Cys80; and Cys49, Cys70, Cys73, Cys76) are predicted to coordinte two [4Fe-4S] clusters (Figure S10). The distance between the two [4Fe-4S] clusters is approximately 10 Å, and the distance between one [4Fe–4S] cluster and the heme iron center in FsnU2 is around 12 Å, both within the range compatible with efficient electron transfer (Figure S11).

**Figure 4.**
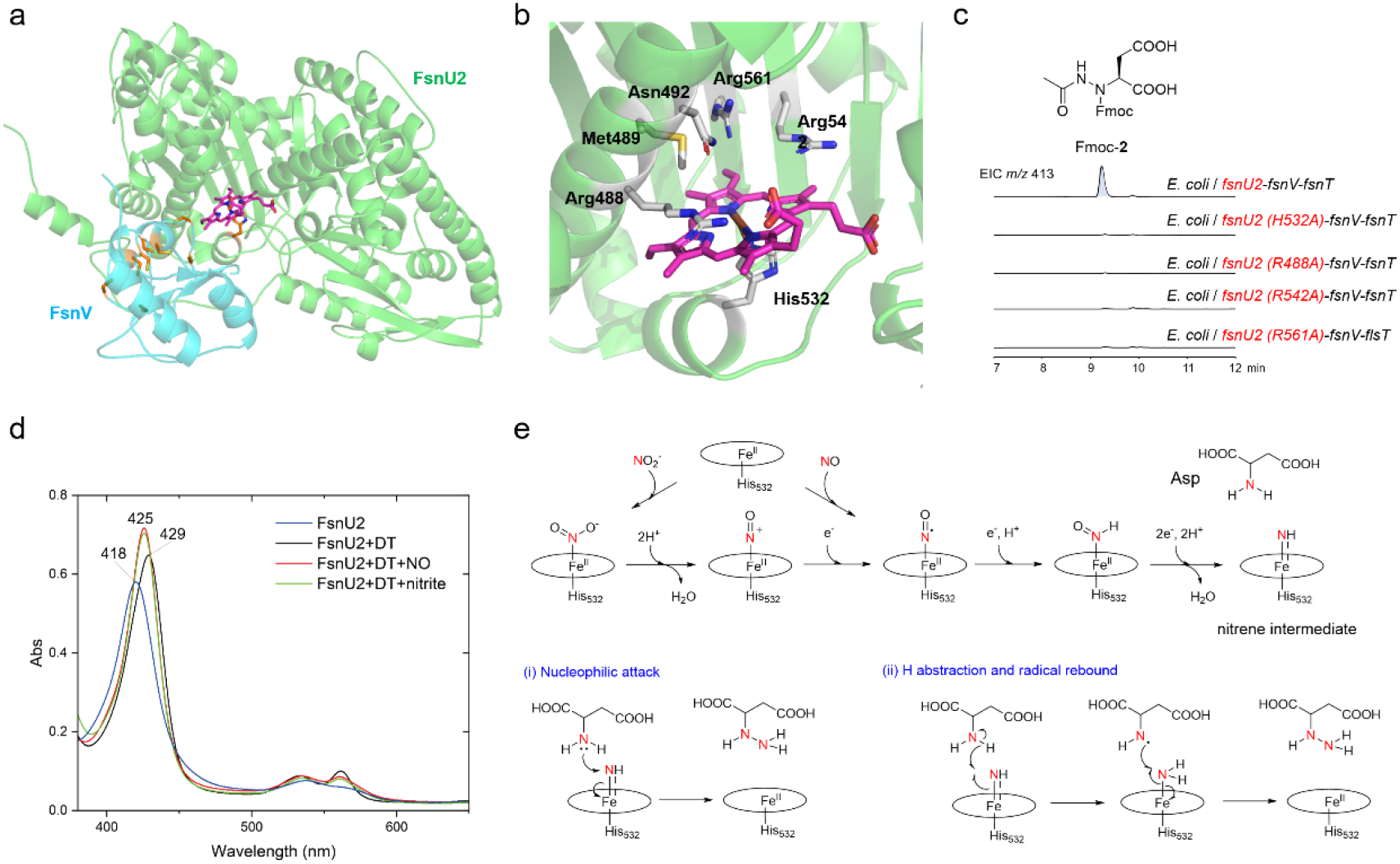
Mechanistic insights into hydrazinosuccinic acid biosynthesis. (a) The AlphaFold-predicted structural model of FsnU2-FsnV complex. The heme is shown in magenta, and residues predicted for heme coordination and 4Fe-4S cluster formation are indicated as orange stick. (d) The predicted active site of FsnU2. The conserved residues are indicated as light grey stick. (c) Analysis the roles of conserved residues (H532A, R488A, R542A, and R561A) using in vivo reconstitution. (d) UV-Vis spectra of dithionite (DT)-reduced FsnU2 incubated with nitrite or NO (provided using the NO donor Diethylamine NONOate). (e) Proposed reaction mechanism for N-N bond formation catalyzed by the FsnU2-FsnV complex.

To assess the functional importance of the heme-coordinating histidine residue in FsnU2, we performed site-directed mutagenesis of His532. The resulting H532A variant was unable to produce compound **2** in the in vivo bioconversion assay (Figure 4c). We further observed that several residues on the distal side of the heme cofactor are highly conserved, suggesting their involvement in catalysis or substrate binding. Among these, three invariant arginine residues (Arg488, Arg542, and Arg561) were individually mutated to alanine. Each of the resulting variants (R488A, R542A, and R561A) completely lost the ability to generate compound **2**, highlighting their essential roles in enzymatic function (Figure 4c). Additionally, UV–visible spectroscopic analysis revealed a shift in the Soret band of dithionite-reduced FsnU2 from 429 to 425 nm upon the addition of nitrite or an NO donor, supporting the direct binding of nitrogen oxide species to the heme iron (Figure 4d). Together, these results strongly support the conclusion that the heme cofactor in FsnU2 plays a direct catalytic role in the N–N bond-forming reaction. Based on these results, we propose a mechanistic model for the FsnU2/FsnV enzyme pair (see Discussion) (Figure 4e).

## Discussion

Enzymes that catalyze N–N bond formation have garnered increasing attention due to their unique chemistries and important roles in both primary and secondary metabolism. Several such enzymes with distinct catalytic mechanisms have been identified to date [28–31]. One of the examples is the heme-dependent hydrazine synthase involved in the microbial nitrogen cycle. This enzyme catalyzes the formation of hydrazine (N2H4) from ammonia and nitric oxide, playing a central role in the anammox (anaerobic ammonium oxidation) pathway [32,33]. In the field of natural product biosynthesis, piperazate synthases such as KtzT have been shown to catalyze the intramolecular N–N bond formation leading to the cyclization of ^5^*N*-hydroxy-ornithine to yield piperazate, a rare non-proteinogenic amino acid found in several bioactive compounds [23,34]. Beyond heme enzymes, ATP-dependent N–N bond-forming enzymes have also been identified, expanding the mechanistic diversity of N–N bond-forming biocatalysts [9–11,35–37]. Collectively, these discoveries highlight the remarkable catalytic repertoire of enzymes involved in N–N bond formation.

Unlike all previous reported examples, the FsnU2/FsnV complex appears to catalyze an intermolecular N–N bond formation between an inorganic nitrogen oxide species (e.g., nitrite or nitric oxide) and an amino acid substrate in a heme-dependent manner. To couple NO or NO_2_^−^ with the α-amine of amino acid substrate to form a hydrazine product, the nitrogen oxide species must be reduced. This requirement is consistent with the essential role of the cognate [4Fe-4S] ferredoxin partner FsnV, both in vivo and in vitro, which potentially transfers electrons to the heme iron center of FsnU2 to facilitate the reductive activation of NO_x_ species. Conserved polar residues on the distal side of the heme cofactor may play key roles in proton transfer or substrate recognition. Building on the mechanistic insights from the heme-dependent enzymes including hydrazine synthase, piperazate synthase and nitrite reductase, we propose a catalytic mechanism for FsnU2/FsnV-catalyzed N-N bond formation (Figure 4e). In this model, the ferrous heme iron in FsnU2 coordinates to NO or NO_2_− through the nitrogen atom. Following electron transfer from FsnV, the bound nitrogen oxide is reduced to a reactive intermediate, possibly a nitrene species. Such nitrene intermediates have recently been implicated in piperazate synthase-catalyzed N–N formation [34], and in engineered heme enzyme-catalyzed hydroxylamine amination reactions [38]. Meanwhile, the α-amine of L-Asp is positioned in close proximity to this nitrene intermediate, allowing for N-N bond formation through either nucleophilic attack or a hydrogen abstraction followed by radical rebound mechanism. It is worth noting that, while this manuscript was in preparation, a related heme-dependent enzyme named NegJ, from the biosynthetic pathway of antibiotic negamycin, was reported to catalyze a similar transformation using nitrite and glycine as substrates [39]. Additionally, a preprint was recently posted describing in vitro characterization of KinJ/KinI enzyme pair [40]. Together, these results unveil a new family of heme-dependent enzymes that catalyze intermolecular N–N bond formation using inorganic nitrogen oxide species. Despite these findings, the precise catalytic mechanism and structural basis of this emerging enzyme family remain poorly understood. Future structural and spectroscopic studies will be crucial for fully understand their reaction mechanisms. In silico genome mining has revealed that FsnU2 homologs are widely distributed across bacterial genomes, suggesting that this protein family may serve as a powerful genetic marker for discovering novel natural products bearing N–N motifs (Figure S13).

In conclusion, we have characterized a new heme-dependent enzyme system that catalyzes an intermolecular N–N bond formation between Asp and nitrite or nitric oxide to produce hydrazinosuccinic acid. This work not only expands the current enzymatic toolkit for N–N bond formation but also sheds light on a novel enzymatic strategy for nitrogen incorporation. Our findings lay the foundation for future genome mining and synthetic biology applications aimed at harnessing N–N biocatalysts for production of nitrogen-containing compounds.

## Acknowledgments

This work was supported by the National Key R&D Program of China (2025YFA0921001) and the National Natural Science Foundation of China (32350051) to Y.-L. Du, and China Postdoctoral Science Foundation (2024M752836, 2024T170784) and Postdoctoral Fellowship Program of CPSF (GZB20230658) to G. Zhao.

