## Supporting Information for "Genome Mining and Characterization of a Heme-Dependent Enzyme Catalyzing Intermolecular Nitrogen–Nitrogen Bond Formation in Hydrazinosuccinic Acid Biosynthesis"

### Materials and methods

#### General note

DNA primiers were purchased from Tsingke Biological Technology. Synthetic genes were purchased from Tsingke Biological Technology. Reagents were purchased from suppliers including Sigma-Aldrich, Thermo Fisher Scientific, Cambridge Isotope Laboratories, Aladdin, Sangon Biotech, Bio Basic Inc.

DNA manipulations in *Escherichia coli* and *Streptomyces* were carried out according to standard procedures [1,2]. Selection of recombinant *E. coli* and *Streptomyces* strains was carried out using kanamycin (50 μg/mL), ampicillin (100 μg/mL), spectinomycin (50 μg/mL), thiostrepton (50 μg/mL) and nalidixic acid (25 μg/mL).

#### The analysis of metabolites by LC-MS

LC-MS analysis was performed using an Agilent 1260 II-6125 system with an Agilent EC-C18 column (4 μm, 4.6 mm ID× 150 mm). There are two methods for analysis. The first analytic method was for identifying products from in vivo biotransformation and in vitro biochemical assays. Elution used a flow rate of 0.8 mL/min with a linear gradient of water:acetonitrile (75:25 to 5:95, v/v) over 20 min. The second analytic method was used to detect albofungin from the fermentation products of engineered *Streptomyces*. The chromatographic separation employed a constant flow rate of 1 mL/min with the following gradient profile: initial isocratic elution at water:acetonitrile (90:10) for 3 minutes, followed by a linear gradient of water:acetonitrile (90:10 to 10:90, v/v) over 15 minutes, and concluding with isocratic elution at 100% acetonitrile for 5 minutes. Both components of the mobile phase contained 0.05% (v/v) formic acid.

#### Construction of engineered *E. coli* strains

The pCDFDuet-*fsnT*-*fsnU2* plasmid was constructed as follows: The *fsnT* coding region was amplified from *Micromonospora* sp. JCM 30529 genomic DNA using 2×Phanta Max Master Mix (Vazyme), digested with NcoI and HindIII, and ligated into the corresponding sites of pCDFDuet-1. Subsequently, the *fsnU2* coding sequence was amplified from *Micromonospora* sp. JCM 30529, digested with NdeI and XhoI, and ligated into the corresponding sites of the resulting pCDFDuet-*fsnT* plasmid. For the creation of pCDFDuet-*fsnT*-*kinJ* plasmid, the *kinJ* coding region was amplified from synthetic gene, digested with NdeI and XhoI, and ligated into the respective site of pCDFDuet-*fsnT*.

To construct the pCOLADuet-*fsnV* plasmid, the *fsnV* coding region was amplified from *Micromonospora* sp. JCM 30529 genomic DNA using 2×Phanta Max Master Mix, Vazyme), digested with NcoI and XhoI, and cloned into the corresponding sites of pCOLADuet-1 to generate the pCOLADuet-*fsnV* plasmid.

For the construction of the pCDFDuet-*fsnT*-*kinJ* plasmid, the *kinJ* coding region was amplified from synthetic gene. Following NdeI and XhoI digestion, the resulting PCR product were ligated into the corresponding site within pCDFDuet-*fsnT* vector.

For the creation of the pCOLADuet-*kinI* plasmid, the *kinI* coding region was amplified from synthetic gene, digested with NcoI and XhoI, the resulting PCR product was ligated into the respective site within pCOLADuet-1 vector.

For the construction of pCDFDuet-*fsnT*-*fsnU2* point mutants, site-directed mutagenesis of pCDFDuet-*fsnT*-*fsnU2* was performed using the pCDFDuet-*fsnT*-*fsnU2* as a template, amplified with complementary primers harboring target mutations and 2×phanta Max Master Mix. After DpnI-mediated template elimination, the reaction mixture was transformed into *E. coli* DH5α competent cells and the correctness of mutant plasmids was verified by DNA sequencing.

**Biotransformation and metabolic analysis of engineered *E. coli* strains**

For the bioconversion for engineered *E. coli* strains, cultures were grown at 37 °C and 180 rpm in Luria broth (LB) medium supplemented with the appropriate antibiotic until the $OD_{600}$ reached approximately 0.4-0.6. Protein expression was induced by the addition of 0.1 mM isopropyl β-D-1-thiogalactopyranoside (IPTG). Subsequently, the cells were harvested via centrifugation and resuspended in M9 medium containing 0.4% glucose, 2 mM $MgSO_4$, 0.1 mM $CaCl_2$, 0.1% L-aspartate and 3 mM $NaNO_2^-$(or isotopically labeled $^{15}N$-L-aspartate and $^{15}N$-$NaNO_2^-$ for precursor feeding experiments). After incubation at 30°C and 180 rpm for 12 hours, the cultures were centrifuged and the supernatant was collected. To precipitate proteins, nine volumes of methanol were added to the supernatant and maintained at -20°C for 30 minutes. The samples were then centrifuged, and the supernatant volumes were reduced under vacuum. For LC-MS analysis, samples were centrifuged once more, and 8 μL of the resulting supernatant was utilized for LC-MS analysis.

**Cloning, expression, purification and analysis of recombinant proteins**

For the purification of FsnT protein, the *fsnT* coding region was amplified from

*Micromonospora* sp. JCM 30529 genome using 2×Phanta Max Master Mix (Vazyme) with primer 28a-*fsnT*-NdeI-F and 28a-*fsnT*-XhoI-R. Following NdeI and XhoI digestion, the resulting PCR product was ligated into the corresponding site within the pET28a vector to generate pET28a-FsnT plasmid. After sequence confirmation, the plasmid was transformed into *E. coli* for protein expression. The cultures were grown in Luria broth (LB) medium with the appropriate antibiotic at 37 °C, 180 rpm until the $OD_{600}$ reached 0.4-0.6. The protein expression was induced by the addition of 0.1 mM IPTG at 18 °C for 18 h. Subsequently, the cells were harvested by centrifugation (5000 rpm, 8 min), resuspended in 25 mL lysis buffer (25 mM Hepes, 300 mM NaCl and 10 mM imidazole, pH 8.0) and lysed by sonication on ice. After centrifugation (15,000 rpm, 40 min, 4°C), the supernatant was loaded onto Ni-NTA resin. After being washed with washing buffer (300 mM NaCl, 25 mM Hepes, 30 mM imidazole, pH 8.0), the proteins of interest were eluted with elution buffer (300 mM NaCl, 25 mM Hepes, 50 mM or 300 mM imidazole, pH 8.0). The proteins were analyzed by SDS-PAGE and dialyzed against the storage buffer containing 25 mM Hepes, 100 mM NaCl, 5% glycerol (pH 8.0). The dialyzed proteins were concentrated and stored at -80 °C for further use. Protein concentrations were determined spectrophotometrically at 280 nm using extinction coefficients calculated via ExPASy ProtParam.

For the purification of FsnU2, the *fsnU2* coding region was amplified from the *Micromonospora* sp. JCM 30529 genome using Phanta Max Master Mix (Vazyme) with primers 28a-*fsnU2*-NdeI-F and 28a-*fsnU2*-XhoI-R. The PCR product was digested with NdeI and XhoI and ligated into the corresponding sites of the pET28a vector, generating plasmid pET28a-*fsnU2*. Following sequence verification, this recombinant plasmid was transformed into *E. coli* for expression. Cells were grown at 37°C and 180 rpm in LB medium supplemented with the appropriate antibiotic until $OD_{600}$ reached 0.4-0.6. Protein expression was induced by adding 1 mM ammonium iron(II) sulfate, 1 mM 5-aminolevulinic acid, and 0.1 mM IPTG. After incubation at 18°C for 18 h, cells were harvested by centrifugation (5000 rpm, 8 min), resuspended in lysis buffer (25 mL; 25 mM Tris-HCl, 300 mM NaCl, 10 mM imidazole, pH 8.0), and lysed by sonication on ice. Lysates were clarified by centrifugation (15,000 rpm, 40 min, 4°C) to separate insoluble fractions. His-tagged proteins were purified from the soluble fraction using Ni-NTA affinity chromatography. The resin was washed with washing buffer (25 mM Tris-HCl, 300 mM NaCl, 30 or 50 mM imidazole, pH 8.0), and bound proteins were eluted with elution buffer (25 mM Tris-HCl,

300 mM NaCl, 300 mM imidazole, pH 8.0). Eluted proteins were analyzed by SDS-PAGE, dialyzed into storage buffer (50 mM Tris-HCl, 100 mM NaCl, 10% glycerol, pH 8.0), concentrated, aliquoted, and stored at -80°C. Protein concentration was determined using a Bradford assay kit.

For the purification of FsnV protein, the *fsnV* coding region were amplified from *Micromonospora* sp. JCM 30529 genomic DNA using 2×Phanta Max Master Mix (Vazyme) with primer 22b-*fsnV*-NdeI-F and 22b-*fsnV*-XhoI-R. Following NdeI and XhoI digestion, the resulting PCR product was cloned into the corresponding sites of pET22b, yielding pET22b-*fsnV* plasmid. After sequence verification, the plasmid was transformed into *E. coli* for protein expression. The cultures were grown at 37 ℃ and 180 rpm in Luria broth (LB) medium with the appropriate antibiotic to an $OD_{600}$ of 0.4-0.6. Protein expression was induced by the addition of 1 mM ammonium iron(II) sulfate, 100 μM cysteine and 0.1 mM IPTG, followed by 18-h incubation at 18°C. Cells were harvested by centrifugation at 5000 rpm for 8 min and resuspended in 25 mL lysis buffer (25 mM Tris-HCl, 300 mM NaCl and 10 mM imidazole, pH 8.0). Subsequently, the cells were sonicated on ice and centrifuged (4 ℃, 15000 rpm, 40 min) to separate soluble and insoluble fractions. Clarified lysate was loaded onto Ni-NTA resin. After being washed with washing buffer (300 mM NaCl, 25 mM Tris-HCl, 30 mM or 50 mM Imidazole, pH 8.0), the protein of interest were eluted with elution buffer (300 mM NaCl, 25 mM Tris-HCl, 300 mM imidazole, pH 8.0). The proteins were analyzed by SDS-PAGE and dialyzed twice into the storage buffer containing 50 mM Tris-HCl, 100 mM NaCl, 10% glycerol (pH 8.0). The dialyzed proteins were concentrated and stored at -80 ℃ for further use. Protein concentration was determined by Bradford assay kit.

For the purification of FsnU2 and FsnV coexpression protein, the *fsnV* coding regions were amplified from *Micromonospora* sp. JCM 30529 genome using 2×Phanta Max Master Mix (Vazyme) with primer *fsnV*-NcoI-F and *fsnV*-HindIII-R. The digested fragment was ligated into pCDFDuet-1 vector to generate pCDFDuet-*fsnV* plasmid. After sequencing verification, this plasmid and pET28a-*fsnU2* were co-transformed into *E. coli* for protein expression. As a control, pET28a-*fsnU2* and pCDFDuet-1 were co-transformed into *E. coli*. The cultures were grown at 37 ℃ and 180 rpm in Luria broth (LB) medium with the appropriate antibiotic until the $OD_{600}$ reached 0.4-0.6. Protein co-expression was induced by the addition of 1 mM ammonium iron (II) sulfate, 1 mM 5-aminolevulinic acid, 100 μM cysteine (except for the FsnU2-pCDFDuet co-expression strain) and 0.1 mM IPTG, followed by 18 h incubation at 18°C. Cells were harvested via centrifugation at

5000 rpm for 8 min, resuspended in 25 mL lysis buffer (25 mM Tris-HCl, 300 mM NaCl and 10 mM imidazole, pH 8.0), and lysed by sonication on ice. The lysate was clarified by centrifugation at 15,000 rpm, 4°C for 40 min, and the supernatant was loaded onto Ni-NTA resin. After washing with buffer containing 25 mM Tris-HCl, 300 mM NaCl, and 30 mM or 50 mM imidazole (pH 8.0), the proteins of interest were eluted with elution buffer (300 mM NaCl, 25 mM Tris-HCl, 300 mM imidazole, pH 8.0). The proteins were analyzed by SDS-PAGE and dialyzed into the storage buffer containing 50 mM Tris-HCl, 100 mM NaCl, 10% glycerol (pH 8.0). Target complexes were confirmed by SDS-PAGE, dialyzed against storage buffer (50 mM Tris-HCl, 100 mM NaCl, 10% glycerol, pH 8.0), concentrated by ultrafiltration, and stored at -80°C. The dialyzed proteins were concentrated and stored at -80 °C for further use. Protein concentration was determined by Bradford assay kit.

**Analysis of the cofactor within FsnU2**

The analytical method was as described above. 100 μL of protein was mixed with 1 mL of acetonitrile and 1 μL of 5 M HCl. After centrifugation at 12,000 rpm for 5 min, the supernatant was concentrated to dryness in vacco, then dissolved in 80 μL of methanol. Following centrifugation, 5 μL was injected for LC-MS analysis.

**Reconstitution of iron-sulfate clusters in FsnV**

For the reconstitution of iron-sulfer clusters in FsnV, a reconstitution buffer containing 50 mM Tris-HCl, 100 mM NaCl, 4 mM DTT and 10% glycerol (pH 8.0) was thoroughly degassed with argon and transferred into an anaerobic glovebox under a $N_2$ atmosphere with $O_2$ levels maintained at <0.05 ppm. Purified FsnV (200 μM) was dialyzed twice against the above solution in the glovebox. A 6-fold molar excess of ferric chloride was added dropwise to the protein solution with continuous stirring. After incubation for 10 min, an equimolar amount of sodium sulfide was slowly introduced. The reaction proceeded for 2 h at ambient temperature before dialysis against fresh reconstitution buffer to remove unbound ions. After incubation for 2 hours, it was dialyzed in the above buffer to remove excess ferric chloride and sodium sulfide. Protein concentration was re-determined by Bradford assay prior to enzymatic assays and UV-vis spectroscopy.

**In vitro biochemical assays**

The FsnT enzymatic reaction mixture (100 μL) contained 20 μM FsnT, 1 mM $MgCl_2$, 1 mM acetyl-CoA, and 1 mM hydrazinosuccinate in 50 mM HEPES buffer (pH 8.0). After 2 h incubation at 30°C, reactions were quenched by adding two volumes of acetonitrile. For subsequent Fmoc chloride (Fmoc-Cl) derivatization, 150 μL of quenched solution was mixed with 10 μL of 0.2 M borate buffer (pH 8.2) and 20 μL of 50 mM Fmoc-Cl, then incubated at 25°C for 5 min in the dark. After adding 20 μL of 100 μM amantadine hydrochloride, incubation continued at ambient temperature for 10 min under light protection. Following derivatization, samples were centrifuged, and 8 μL of the supernatant was subjected to LC-MS analysis.

The FsnU2 enzymatic reaction was conducted anaerobically. The reaction mixture (100 μL) for the FsnU2-FsnV coupled assay was prepared in 50 mM Tris-HCl buffer (pH 8.0) containing 30 μM FsnU2, 30 μM FsnV (either as-isolated or reconstituted), 5 mM sodium dithionite, 1 mM L-aspartic acid sodium salt, and either 1 mM sodium nitrite or 1 mM DEA NONOate plus 1 mM hydroxylamine. After incubation at 30 °C for 2 h, reactions were quenched by adding two volumes of acetonitrile. Following Fmoc-Cl derivatization (as described previously) and centrifugation, 8 μL of the supernatant was subjected to LC-MS analysis. For the enzymatic reaction utilizing co-expressed FsnU2 and FsnV proteins, the reaction mixture (100 μL) contained 30 μM of the FsnU2-FsnV complex (co-expressed), 5 mM sodium dithionite, 1 mM L-aspartic acid sodium salt, and either 1 mM sodium nitrite or 1 mM DEA NONOate plus 1 mM hydroxylamine. Following Fmoc-Cl derivatization, the reaction products were analyzed by LC-MS as described previously.

**UV-Vis spectroscopic analysis**

For UV-Vis spectroscopic analysis, FsnU2 was diluted to 4 μM in dialysis buffer to a final volume of 600 μL. Assays were performed in 0.7 mL quartz cuvettes using a Shimadzu UV-2600 spectrophotometer.

For UV-Vis spectroscopic analysis of FsnU2 treated with sodium dithionite, the enzyme was diluted to 4 μM in dialysis buffer, then sodium dithionite was added to a final concentration of 5 mM (final volume 600 μL) in 0.7 mL quartz cuvettes. The spectrum was recorded using a Shimadzu UV-2600 spectrophotometer.

For UV-Vis spectroscopic analysis of FsnU2 incubated with substrates (sodium nitrite or the NO donor DEA NONOate), the enzyme was first diluted to 4 μM in dialysis buffer. Sodium dithionite was then added to a final concentration of 5 mM, followed by substrate addition to a final concentration of 2 mM. The final reaction volume was 600 μL. The Spectra were recorded at 30 °C in 0.7 mL quartz cuvettes using a Shimadzu UV-2600 spectrophotometer.

**Construction and metabolic analysis of engineered *Streptomyces***

For the creaction of *S. albus* DM + *alb* BGC (*Δafn8*)*::kinJ-kinI-fzmRQONML* strain, the *kinJ*/*kinI*-containing gene cassette (*kinJ-kinI-fzmRQONML*) were designed, synthesized (Tsingke Biotech), and digested with NdeI/XhoI and cloned into the integrative vector pYLD20, generating pYLD20-*kinJ-kinI-fzmRQONML*. After sequence verification, the plasmid was passaged through the non-methylating *E. coli* ET12567/pUZ8002 host. Subsequently, it was introduced into *S. albus* DM + *alb* BGC (*Δafn8*) via intergeneric conjugation. Exconjugants exhibiting thiostrepton resistance were validated by PCR to confirm site-specific integration.

The mycelium suspension of *Streptomyces* strains served as the inoculum for 50 mL flasks, each containing 10 mL of tryptic soy broth (TSB) medium. Cultures were incubated with shaking for 24 hours at 30 °C and 180 rpm. Subsequently, 500 µl aliquots of seed cultures were plated on YMG agar. After 4-day static incubation at 30°C, the fermentation product was extracted with ethyl acetate, concentrated to dryness in vacco, dissolved with 2.5 mL of methanol, centrifuged, and 5 µl was used for LC-MS analysis.

**Table S1.** Oligonucleotides used in this study

| Primer | SEQUENCE (5' TO 3') | Usage |
|---|---|---|
| *fsnT*-NcoI-F | AGCAGCCCATGGTGACCTGGAAGACCGAACGGATC | Cloning *fsnT* from *Micromonospora* sp. JCM 30529 genomic DNA to construct pCDFDuet-*fsnT* |
| *fsnT*-HindIII-R | AGCAGCAAGCTTGTCAGGGGTTGAGCACCCAGGC | |
| 28a-*fsnT*-NdeI-F | AGCAGCCATATGACCTGGAAGACCGAACGGATC | Cloning *fsnT* from *Micromonospora* sp. JCM 30529 genomic DNA to construct pET28a-*fsnT* |
| 28a-*fsnT*-XhoI-R | AGCAGCCTCGAGTCAGGGGTTGAGCACCCAGGC | |
| 28a-*fsnU2* -NdeI-F | AGCAGCCATATGAGCGCGGGCGCGGAG | Cloning *fsnU2* from *Micromonospora* sp. JCM 30529 genomic DNA to construct pET28a-*fsnU2* |
| 28a-*fsnU2*-NdeI-R | AGCAGCCTCGAGCACGTCCGGCGCCTGCGTTC | |
| *fsnV*-NcoI-F | AGCAGCCCATGGTGAGTCGGAACACAGTTTCACCGGCG | Cloning *fsnV* from *Micromonospora* sp. JCM 30529 genomic DNA to construct pColADuet-*fsnV* and pCDFDuet-*fsnV* |
| *fsnV*-XhoI-R | AGCAGCCTCGAGTCATGCCGGCCGCCCCGCGGTG | |
| 22b-*fsnV*-NdeI-F | AGCAGCCATATGAGTCGGAACACAGTTTCACCGG | Cloning *fsnV* from *Micromonospora* sp. JCM 30529 genomic DNA to construct pET22b-*fsnV* |
| 22b-*fsnV*-XhoI-R | AGCAGCCTCGAGTGCCGGCCGCCCCGCGGTG | |
| *kinJ*-NdeI-F | AGCAGCCATATGACAAGTTCCGCCGAGCGCAACGG | Cloning *kinJ* from synthetic Fzm cluster to construct pCDFDuet-*fsnT*-*kinJ* |
| *kinJ*-XhoI-R | AGCAGCCTCGAGTCACGAGACCTGCTGCGGC | |
| *kinI*-NcoI-F | AGCAGCCCATGGGCATGCCCACGAACACTTCGGACG | Cloning *kinI* from synthetic gene cluster to construct pCDFDuet-*fsnT*-*kinI* |
| *kinI*-XhoI-R | AGCAGCCTCGAGTCATGACACACCCTCAGCGG | |
| *fsnU2*-R488A-F | TCCGCCGCCATGCAGTACAACTCCTGGCACTTC | Site-directed mutagenesis for pCDFDuet-*fsnT*-*fsnU2*-R488A |
| *fsnU2*-R488A-R | CTGCATGGCGGCGGACATCGCCCAGG | |
| *fsnU2*-H532A-F | CACGGGGCGGTGGCGGCGCGGATCCGG | Site-directed mutagenesis for pCDFDuet-*fsnT*- |

| | | |
|---|---|---|
| *fsnU2*-H532A-R | CGCCGCCACCGCCCCGTGGTGGTGCTGGTC | *fsnU2*-H532A |
| *fsnU2*-R542A-F | AGCATCGCGAGCCCGCAAGGG | Site-directed mutagenesis for pCDFDuet-*fsnT*-*fsnU2*-R542A |
| *fsnU2*-R542A-R | CGGGCTCGCGATGCTGAACCGGATCC | |
| *fsnU2*-R561A-F | GACGTCGCCTTGCTGCGCTGCGAC | Site-directed mutagenesis for pCDFDuet-*fsnT*-*fsnU2*-R561A |
| *fsnU2*-R561A-R | CAGCAAGGCGACGTCGACGAACCCGTTG | |

**Table S2.** Plasmids and strains used in this study

| Plasmids | Description | Reference |
|---|---|---|
| pCDFDuet-*fsnT*-*fsnU2* | An expression plasmid of *fsnT* and *fsnU2* | This study |
| pColADuet-*fsnV* | An expression plasmid of *fsnV* | This study |
| pET28a-fsnU2 | An expression plasmid of *fsnU2* | This study |
| pCDFDuet-fsnV | An expression plasmid of *fsnV* | This study |
| pCDFDuet-*fsnT*-*kinJ* | An expression plasmid of *fsnT* and *kinJ* | This study |
| pColADuet-*kinI* | An expression plasmid of *kinI* | This study |
| pCDFDuet-*fsnT*-*fsnU2*-R488A | A site-directed mutagenesis plasmid for fsnU2 for bioconversion | This study |
| pCDFDuet-*fsnT*-*fsnU2*-H532A | A site-directed mutagenesis plasmid for fsnU2 for bioconversion | This study |
| pCDFDuet-*fsnT*-*fsnU2*-R542A | A site-directed mutagenesis plasmid for fsnU2 for bioconversion | This study |
| pCDFDuet-*fsnT*-*fsnU2*-R561A | A site-directed mutagenesis plasmid for fsnU2 for bioconversion | This study |
| Strains | | |
| *E. coli* DH5α | Using as a host for plasmids construction | Laboratory stock |
| *E. coli* Transetta (DE3) | Using as a host for protein expression | Laboratory stock |
| *S. albus* DM + *alb* BGC | *Streptomyces albus* DM harboring albofungin biosynthetic gene cluster | Laboratory stock [3] |
| *S. albus* DM + *alb* BGC (*Δafn8*) | *afn8* knock out *S. albus* DM strain | Laboratory stock |
| *S. albus* DM + *alb* BGC (*Δafn8*) :: *kinJ-kinI-fzmRQONML* | *afn8* knock out *S. albus* DM strain complemented with synthetic gene cassette (*kinJ-kinI-fzmRQONML*) | This study |

**Figure S1**

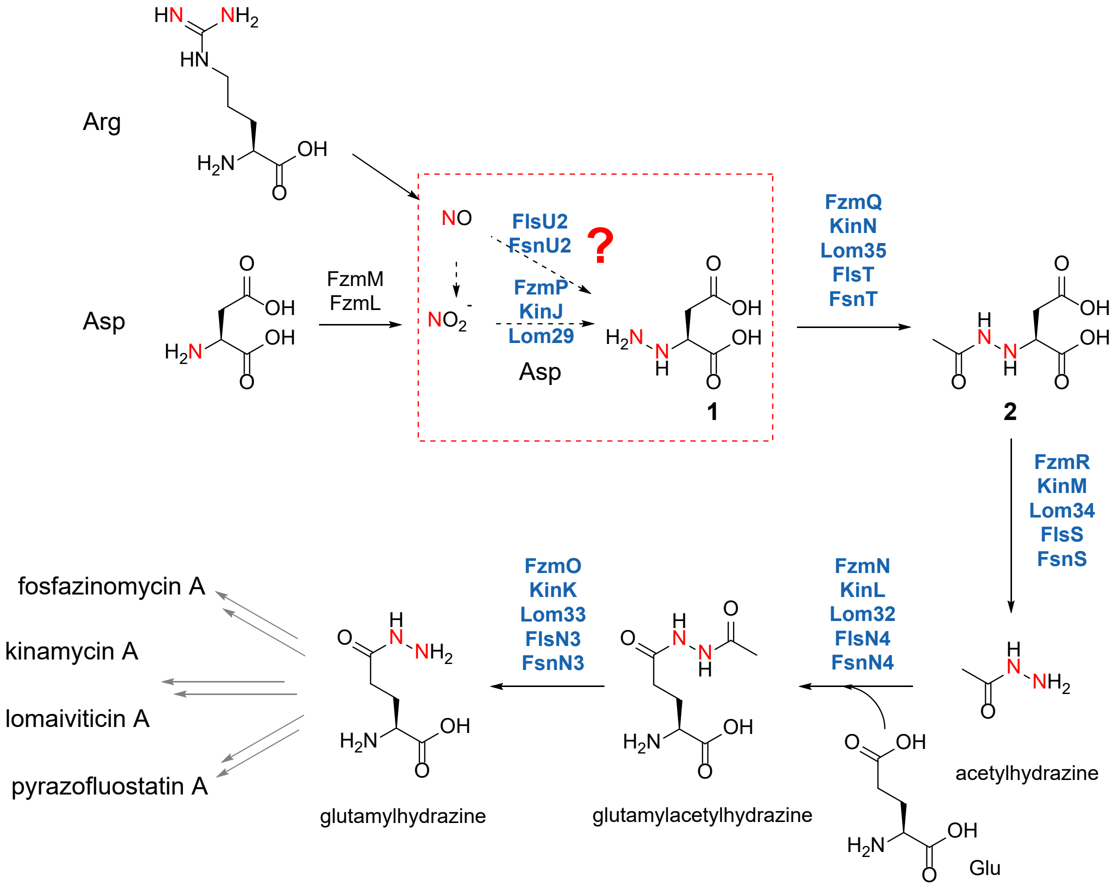


**Figure S1.** The proposed biosynthetic route to glutamylhydrazine [4]. The reaction step investigated by the current study is boxed with red dash line.

### Figure S2

**(a)**

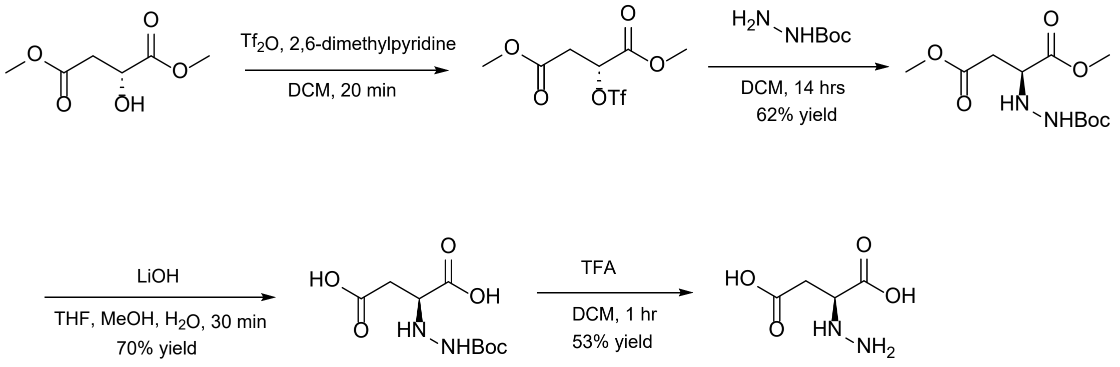
Tf2O, 2,6-dimethylpyridine
DCM, 20 min
H2N NHBoc
DCM, 14 hrs
62% yield
LiOH
THF, MeOH, H2O, 30 min
70% yield
TFA
DCM, 1 hr
53% yield
OH
OTf
HN NHBoc
HO
HN NH2

**(b)**

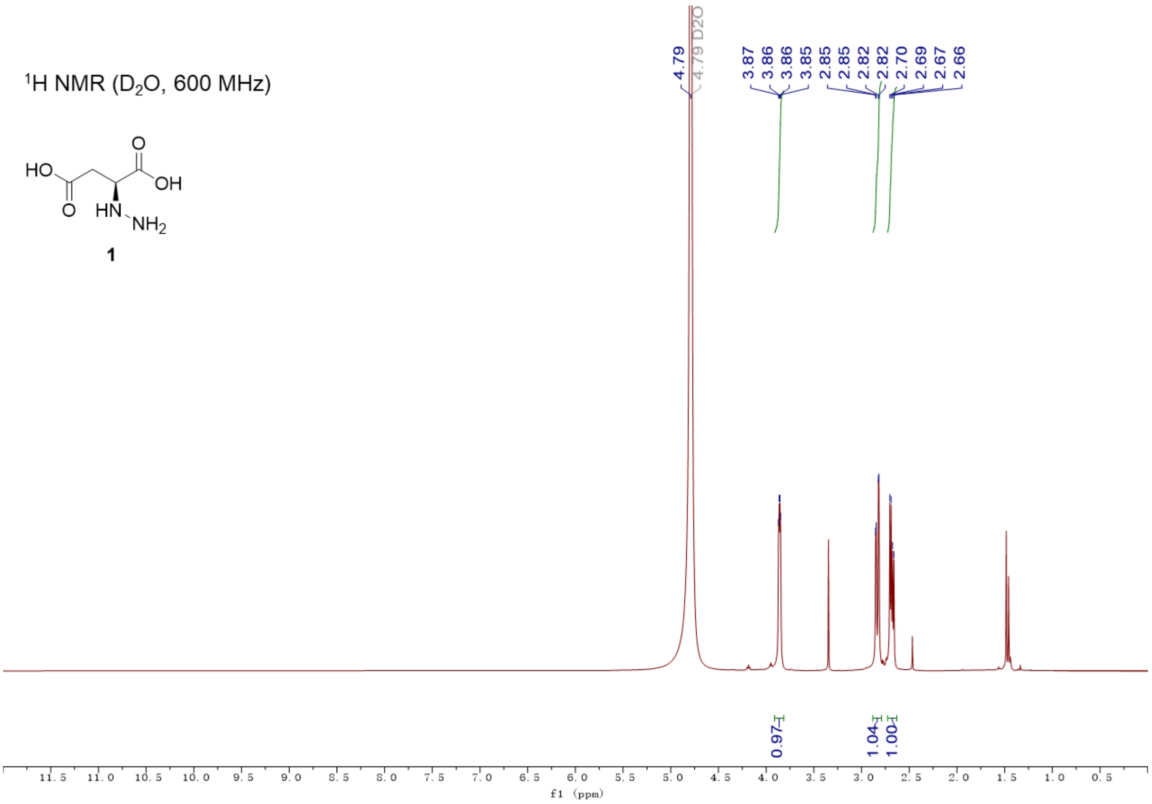
1H NMR (D2O, 600 MHz)
HO
OH
HN NH2
1
4.79
4.79 D2O
3.87
3.86
3.86
3.85
2.85
2.85
2.82
2.82
2.70
2.69
2.67
2.66
0.97
1.04
1.00
f1 (ppm)

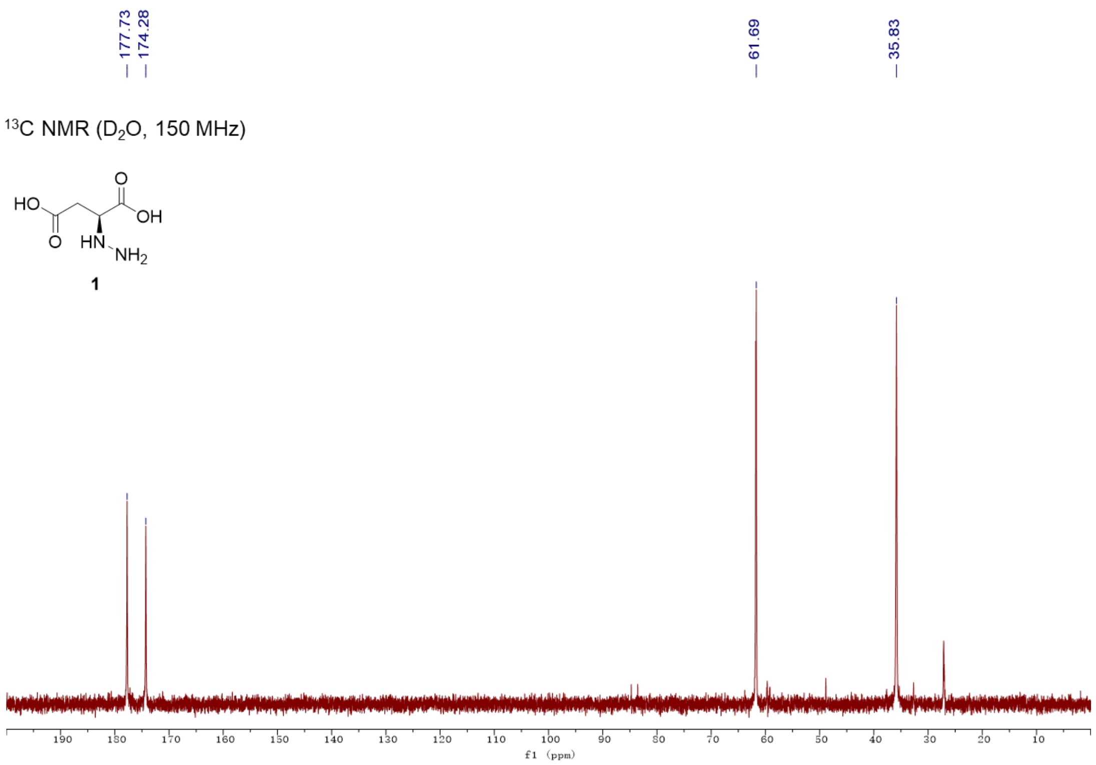


**Figure S2** Chemical synthesis of hydrazinosuccinic acid (**1**). (a) The chemical synthetic route of **1**. (b) $^1$H NMR and $^{13}$C NMR of **1**. $^1$H NMR (500 MHz, $D_2O$): δ 2.68 (dd, 1H, $^2J$ = 17.1 Hz, $^3J$=8.6 Hz), δ 2.84 (dd, 1H, $^2J$ = 17.5 Hz, $^3J$ = 3.4 Hz), δ 3.86 (dd, 1H, $^3J$ = 8.10 Hz, $^3J$ = 3.06 Hz); $^{13}$C NMR (150 MHz, $D_2O$): δ 35.83, δ 61.69, δ 174.28, δ 177.73.

**Figure S3**

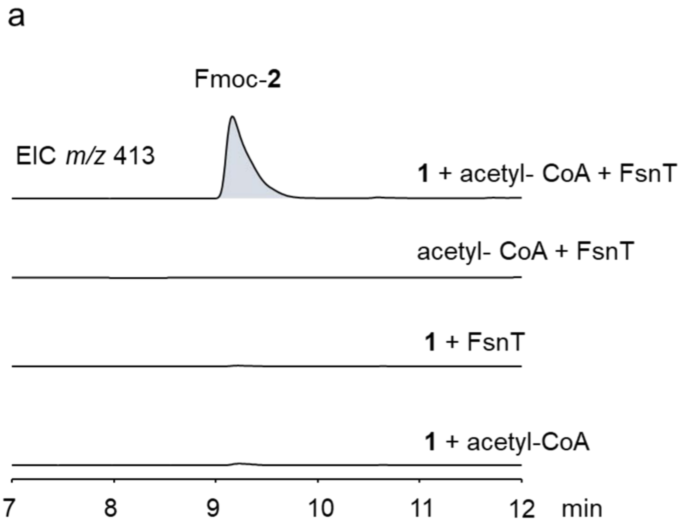


**Figure S3** In vitro biochemical assay of FsnT. (a) In vitro assay of FsnT. EIC = *m/z* 413 for $[M+H]^+$ of Fmoc-**2**. The catalytic activity of FsnT, forming **2** from the substrate and acetyl-CoA, requires the simultaneous presence of the substrate, acetyl-CoA, and the FsnT. (b) SDS-PAGE analysis of His6-FsnT.

**Figure S4**

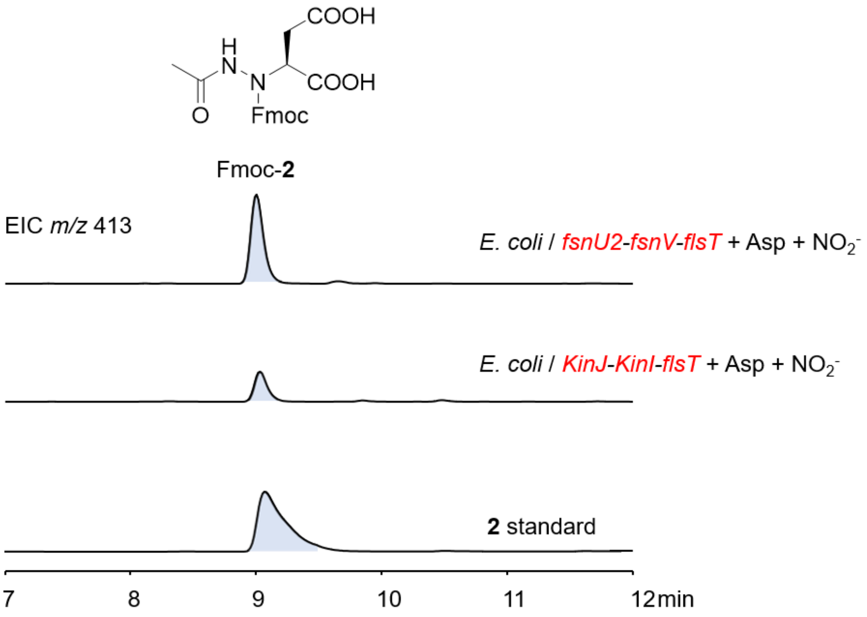


**Figure S4** In vivo biotransformation assay of *kinJ-kinI*. EIC = *m/z* 413 for $[M+H]^+$ of Fmoc-**2**.

**Figure S5**

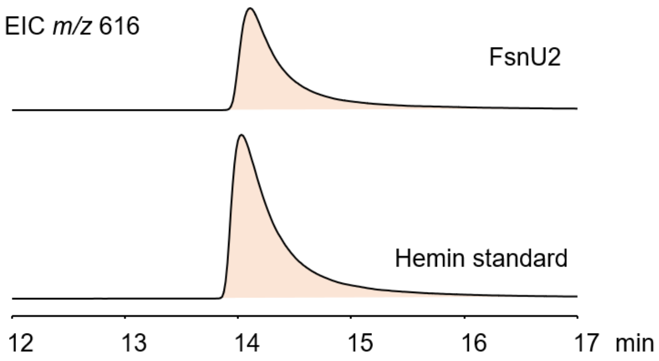


**Figure S5.** LC-MS analysis of the heme b cofactor bound by FsnU2 (EIC = *m/z* 616 for $[M+H]^+$ of hemin).

**Figure S6**

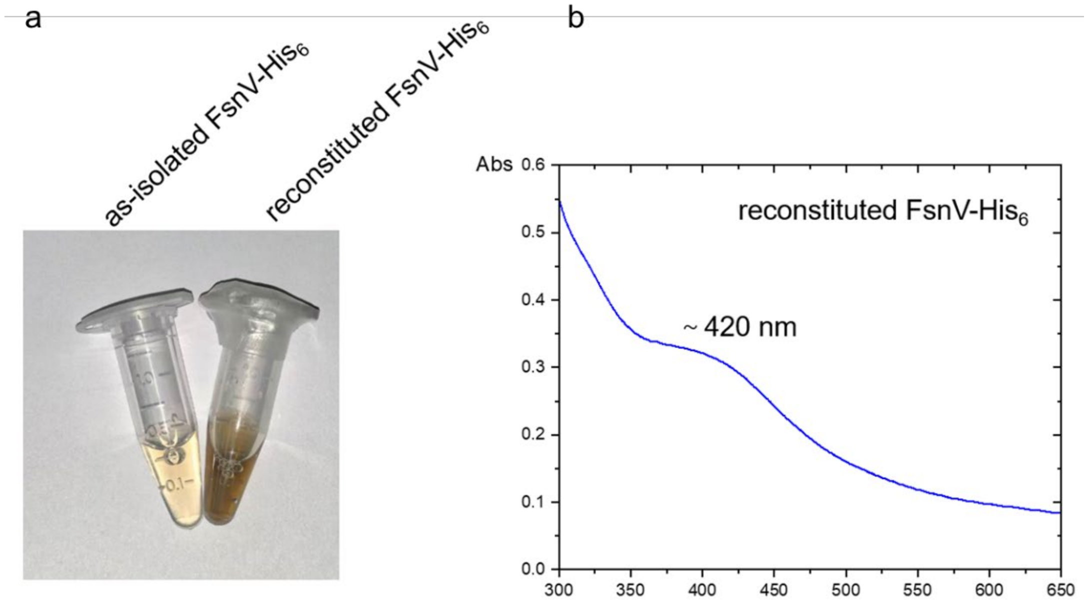


**Figure S6.** Anaerobic reconstitution of the 2[4Fe-4S] ferredoxin FsnV. (a) Color transition of FsnV during iron-sulfur cluster reconstitution. (b) The UV-Vis spectrum of reconstituted FsnV-$His_6$. It has a prominent absorption peak at ~ 420 nm.

**Figure S7**

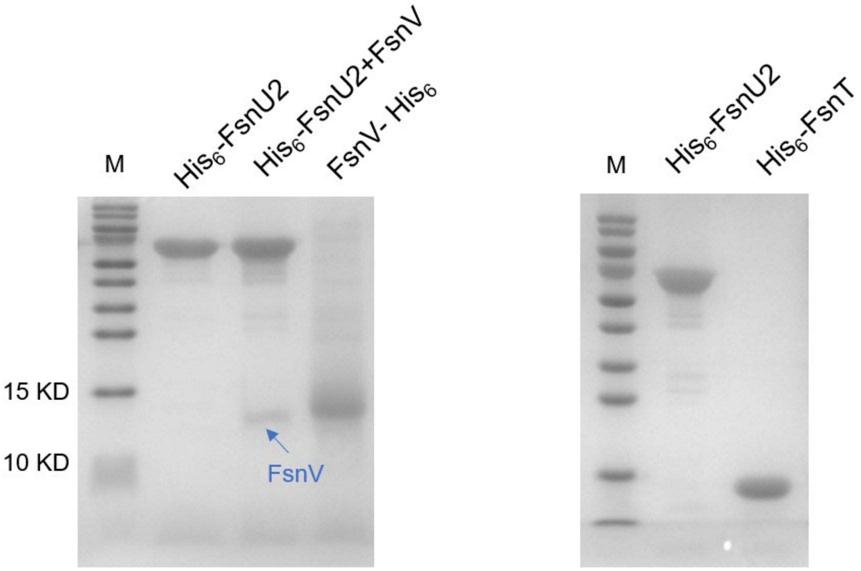


**Figure S7.** SDS-PAGE gels of FsnU2, the complex of $His_6$-FsnU2-FsnV, FsnV-$His_6$, $His_6$-FsnU2 and $His_6$-FsnT. The molecular weight for FsnV, $His_6$-FsnU2 and FsnV-$His_6$ are 12126.79 Da, 70008.16 Da and 13191.91 Da.

**Figure S8**

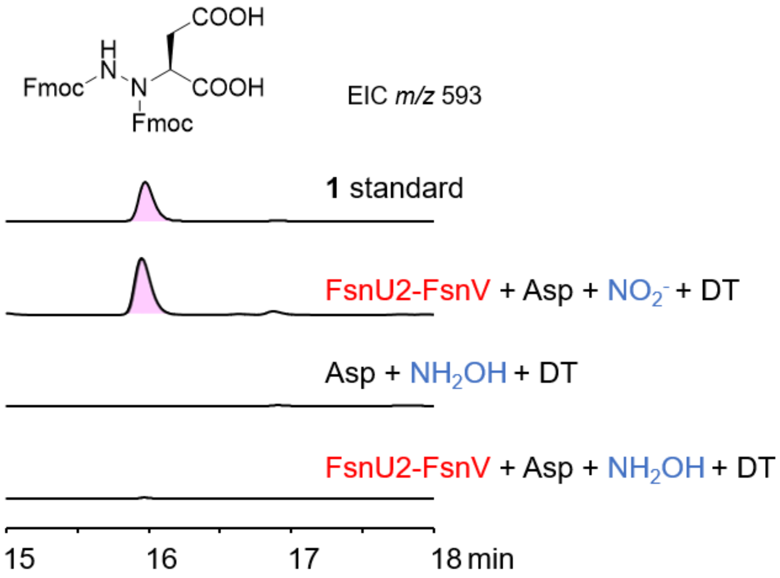


**Figure S8.** In vitro assays of the FsnU2-FsnV using hydroxylamine ($NH_2OH$) as a substrate under anaerobic conditions. The EIC = *m*/*z* 593 ($[M+H]^+$ for bis-Fmoc-**1**) from LC-MS analysis was displayed. The reaction with nitrite as a substrate was used as a positive control.

**Figure S9**

(a)

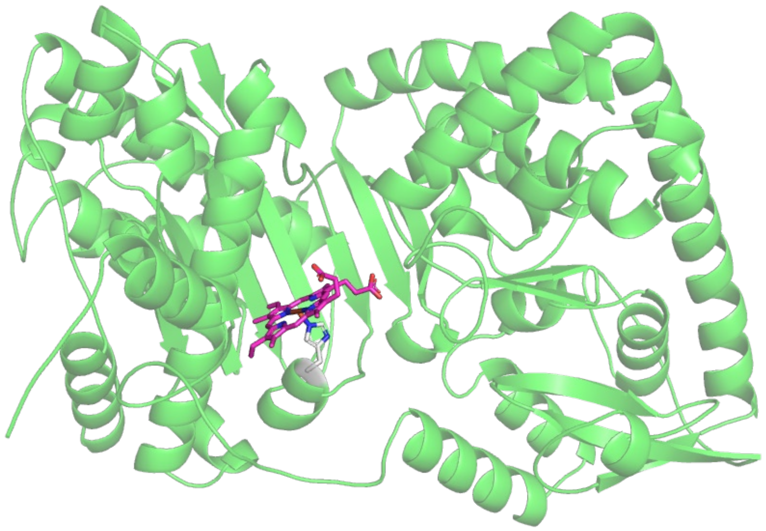

(b)

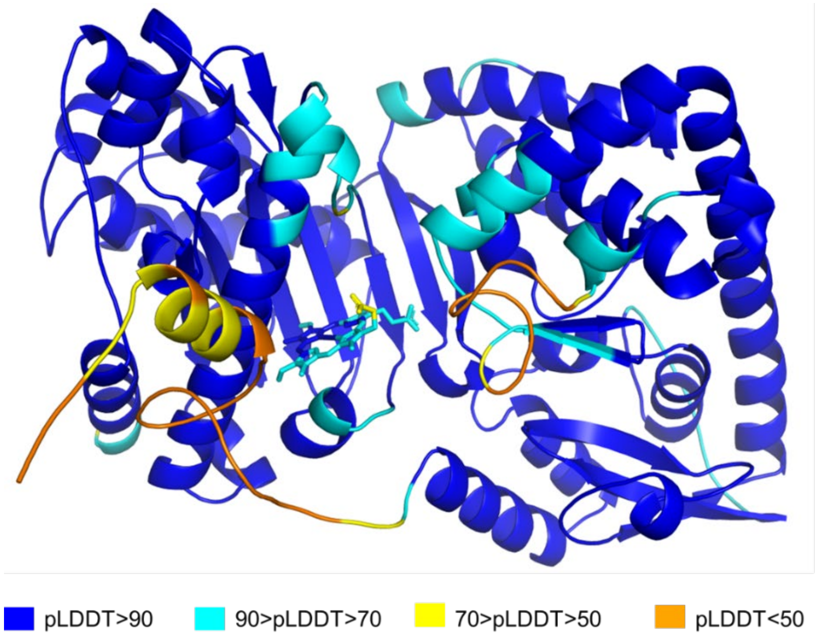


**Figure S9.** The structure of FsnU2 predicted by Alphafold3. (a) The predicted structural model of FsnU2 bound to heme by AlphaFold3. The heme cofactor is shown in magenta and the conserved heme-coordinating His532 is shown in light grey. (b) Predicted Local Distance Difference Test (pLDDT) values for FsnU2 generated by AlphaFold3.

**Figure S10**

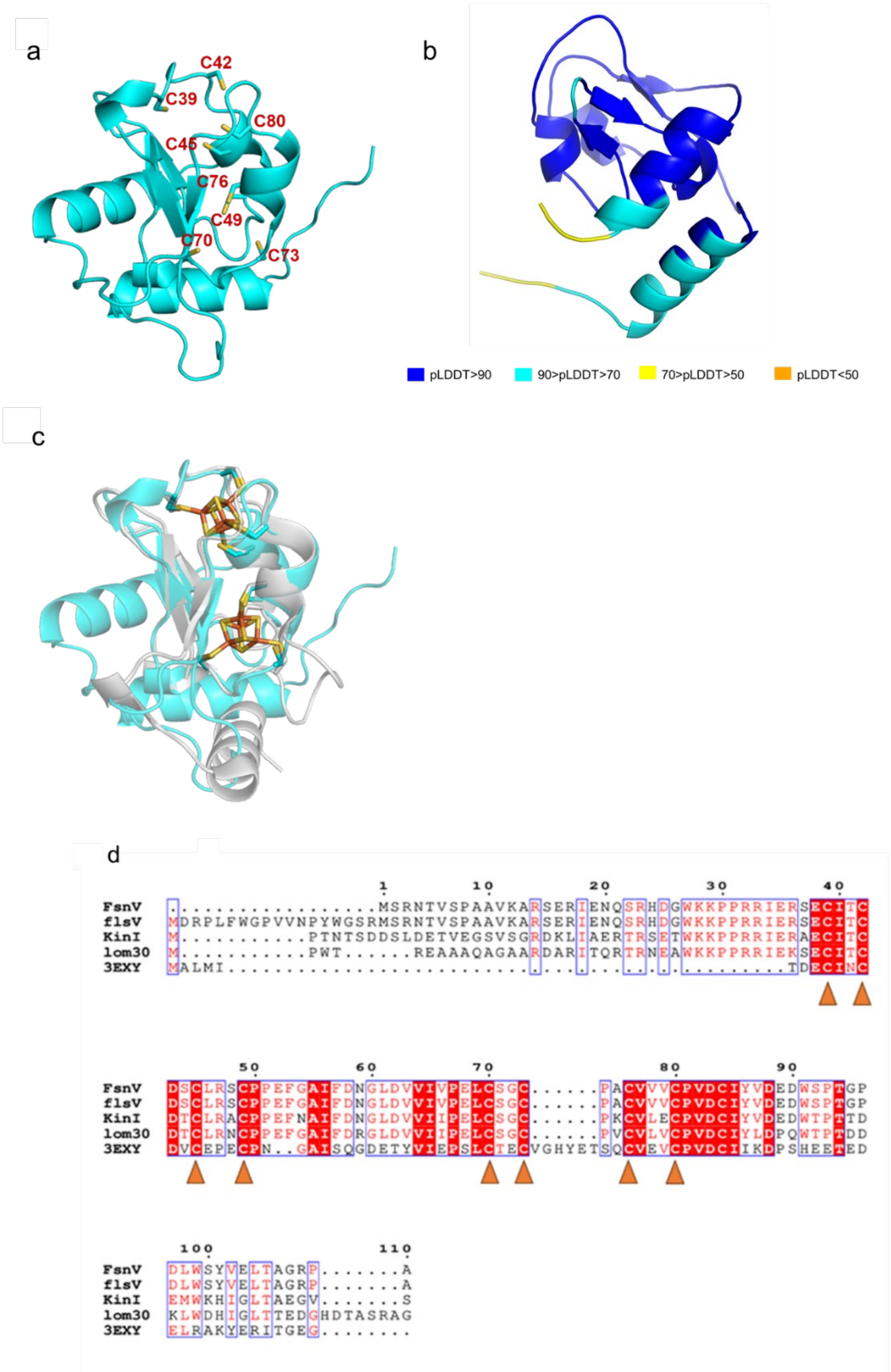


**Figure S10.** The predicted structural model of FsnV and sequence alignment of FsnV and its homologs. (a) The structure of FsnV predicted by alphafold3. (b) Predicted Local Distance Difference Test (pLDDT) values for FsnV generated by AlphaFold3. (c) The structural superposition of FsnV (blue) with its structural homolog, a 2[4Fe-4S] ferredoxin (PDB accession number: 3EXY) from *allochromatium vinosum* (gray) [5]. (d) The multiple sequence alignment of FsnV and its homologs. The conserved residues bonded with iron-sulfur clusters are indicated with orange triangles.

**Figure S11**

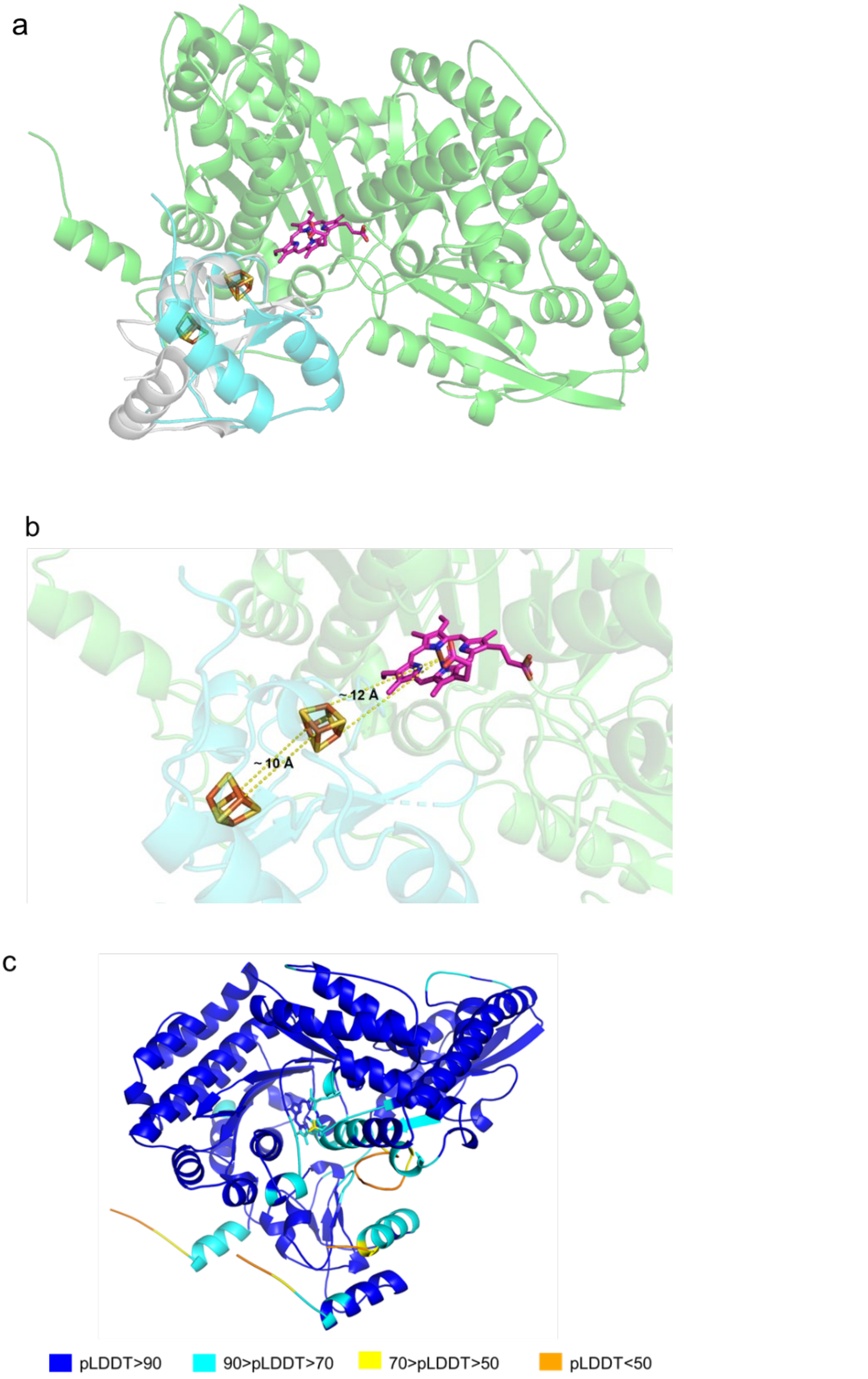


**Figure S11.** The predicted structural model of FsnV-FsnV complex. (a) The structural superposition of the FsnU2-FsnV complex (green and blue) with the ferredoxin crystal structure (3EXY) from *A. vinosum* (gray). (b) Predicted distance between [4Fe-4S] clusters and the heme iron. (c) Predicted Local Distance Difference Test (pLDDT) values for FsnU2-FsnV complex generated by AlphaFold.

**Figure S12**

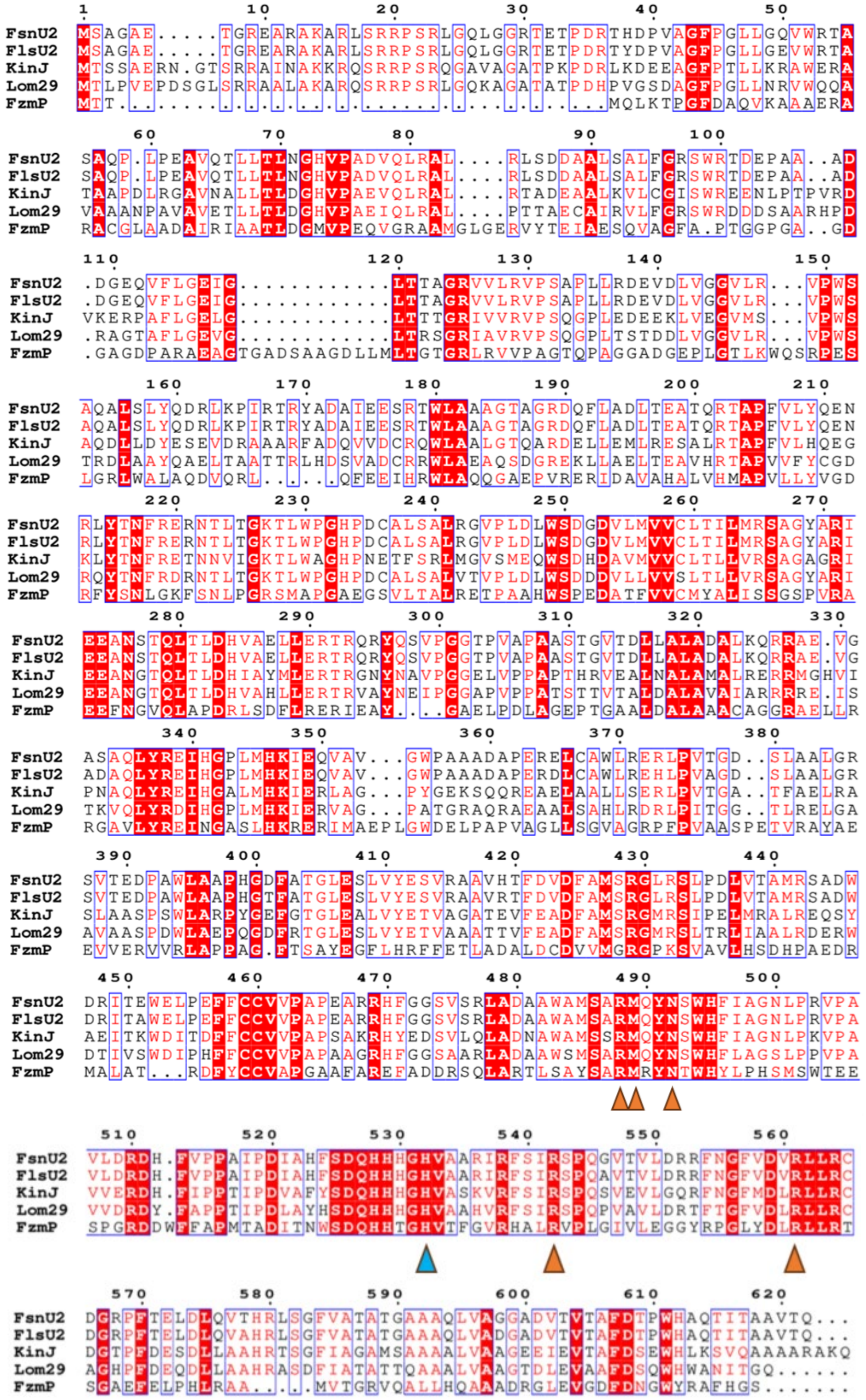

**Figure S12.** Sequence alignment of FsnU2 homologs. The predicted heme-coordinating His residue is indicated with blue triangle. The conserved residues on the distal side of the heme cofactor are indicated with orange triangles.

**Figure S13**

**a**

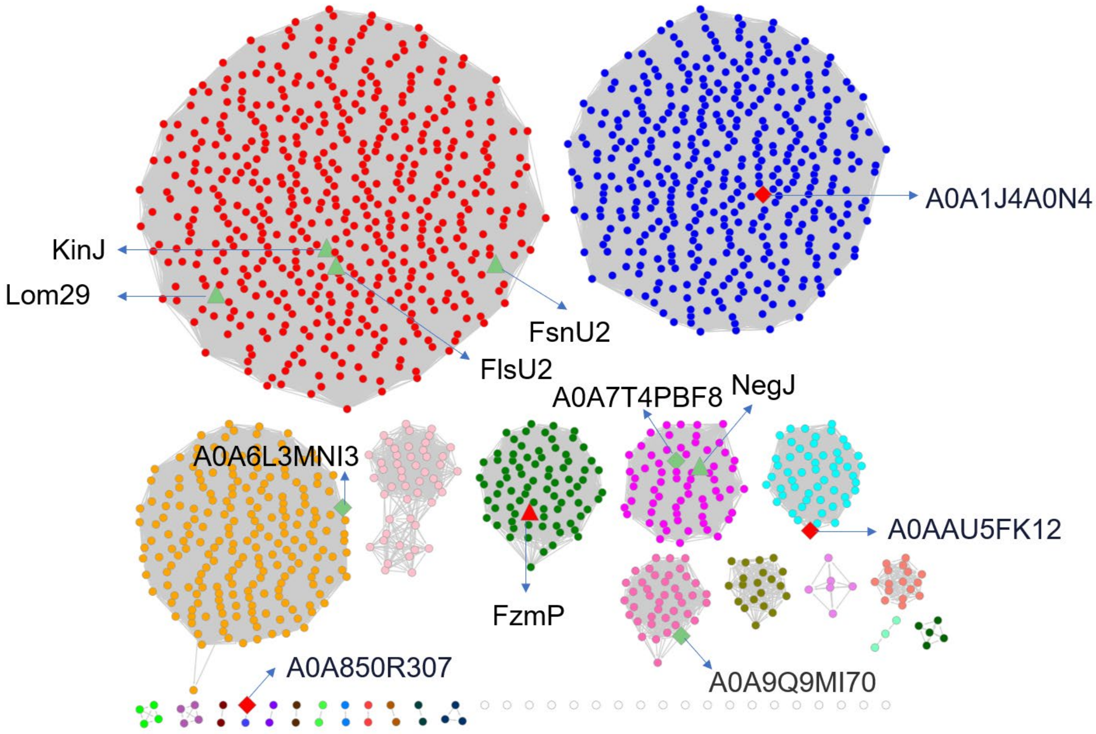

A0A1J4A0N4
KinJ
Lom29
FsnU2
FlsU2
A0A7T4PBF8
NegJ
A0A6L3MNI3
A0AAU5FK12
FzmP
A0A9Q9MI70
A0A850R307

**b**

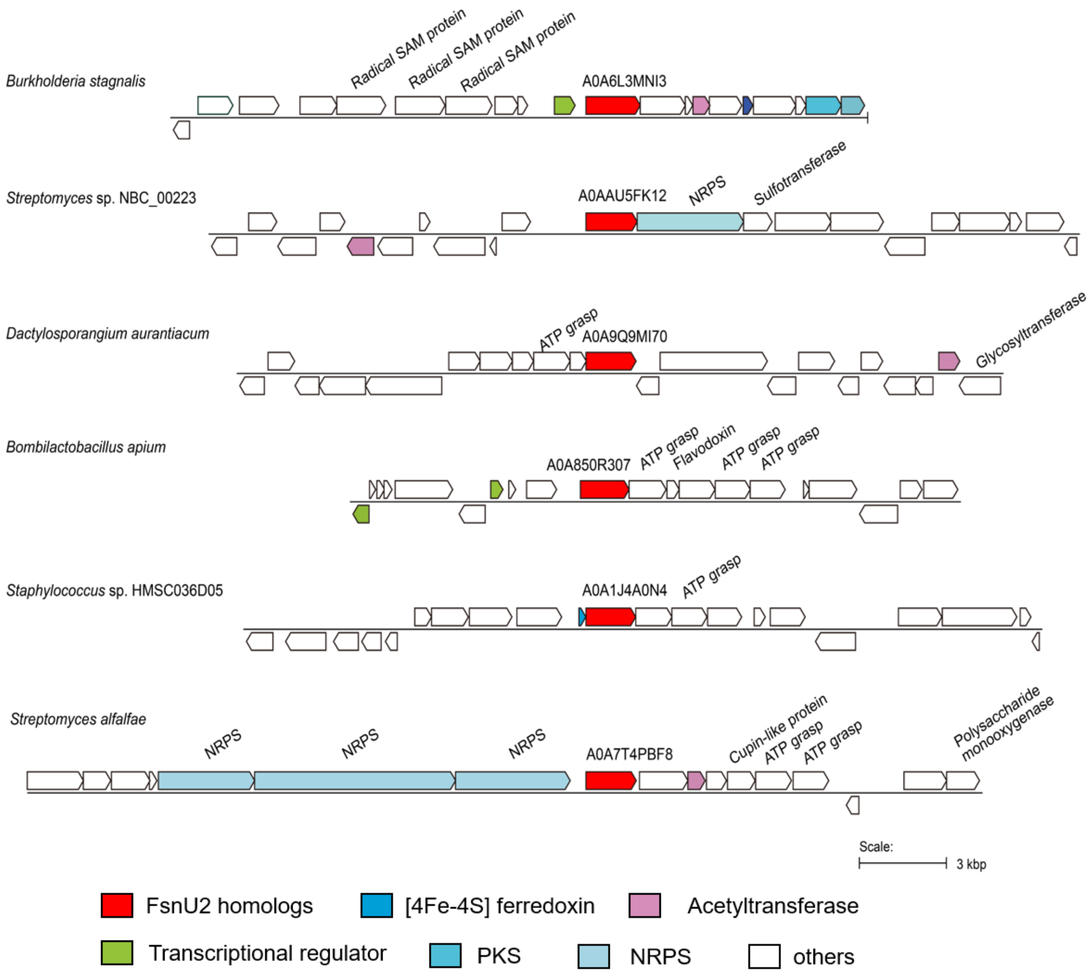


**Figure S13**. In silico mining of putative BGCs using the FsnU2. (a) Sequence similarity network (SSN) analysis of FsnU2 and its homologs. Each node in the network represents a protein and the alignment score is 140. Protein FsnU2, Lom29, KinI, FlsU2, NegJ, FzmP, and the homologs used for Genome neighborhood Tool (GNT) analysis (see panel (b) this figure) are highlighted. (b) Examples of putative FsnU2-containing BGCs obtained using the above SSN and GNT analysis.
